# Genome-wide subcellular protein localisation in the flagellate parasite *Trypanosoma brucei*

**DOI:** 10.1101/2022.06.09.495287

**Authors:** Karen Billington, Clare Halliday, Ross Madden, Philip Dyer, Mark Carrington, Sue Vaughan, Christiane Hertz-Fowler, Samuel Dean, Jack Daniel Sunter, Richard John Wheeler, Keith Gull

## Abstract

*Trypanosoma brucei* is a prototypical trypanosomatid, an important group of human, animal and plant unicellular parasites. Understanding their complex cell architecture and life cycle is hindered since, as with most eukaryotic microbes, ∼50% of the proteins encoded in the genome have completely unknown function. Using fluorescence microscopy and cell lines expressing endogenously tagged proteins we mapped the subcellular localisation of 89% of the proteome, giving clues to function, defining the lineage-specific organelle adaptations for obligate parasitism and mapping the ultra-conserved cellular architecture of eukaryotes. This includes the single flagellum, vital for morphogenesis and pathology: the first comprehensive cartographic analysis of the flagellum in any organism. To demonstrate the power of this resource, we identify novel specialisation of organelle molecular composition through the cell cycle and in specialised subdomains. This is a transformative resource, important for hypothesis generation for both eukaryotic evolutionary molecular cell biology and fundamental parasite cell biology.

## Introduction

The abundance of genome data has transformed molecular cell biology of parasites and model organisms. Although genome sequences have been available for 25 years, there are still many cyptic genes even in the best studied organisms.

Ascribing subcellular localisation of proteins assists understanding function and has largely been addressed through ‘omic’ approaches such as mass spectrometry of purified organelles and hyperplexed organelle localisations by isotope tagging (hyperLOPIT)^1,2^. However, such localisation attributions are limited by the accuracy of organelle purification or fractionation, and sensitivity is limited by protein abundance and characteristics. The gold standard approach for understanding a protein’s sub-cellular localisation and dynamics remains therefore microscopy.

The power of a microscopical map of the subcellular localisation of every protein encoded in an organism’s genome facilitates the localisation to small or rare structures or unpurifiable structures plus cell cycle-dependent changes via the per cell data level. This was a transformative resource for studying *Saccharomyces cerevisiae*^3^ and *Schizosaccharomyces pombe*^4^, and most recently human cell lines^5^. Similar protein positional information in a divergent unicellular parasite would provide powerful diverse opportunities for hypothesis-driven studies.

*Trypanosoma brucei* is a flagellate unicellular parasite, causing sleeping sickness (African trypanosomasis) in humans and nagana in cattle. It is an member of a family of important insect transmitted pathogens, including the human parasites *Leishmania* spp. (leishmaniasis) and *Trypanosoma cruzi* (Chagas disease) and a wide range of animal and plant parasites. *T. brucei* has a complex life cycle alternating between vector and host with multiple developmental forms and adaptations, including characteristic morphologies and specialised surface antigens^6–9^. The flagellum has multiple functions including motility, attachment and environmental sensing^10–12^. However, the flagellum is also a widely conserved organelle in eukaryotes and a defining features of the last eukaryotic common ancestor (LECA), but not yet analysed by genome wide microscopic protein localisation mapping.

*T. brucei* is an early-branching eukaryote (Figure 1A), giving enormous insight into eukaryote evolutionary cell biology and losses/gains in organelle complexity since LECA. It has a highly organised cell with single copies of many organelles (Figure 1B) and a precise division process^13^, allowing unambiguous assignment of cell cycle stages and identification of old and new organelles during and after replication. Protein localisation offers insights into organelle subdomains/dynamics and cell cycle-dependent localisation changes. Here, we generate a high quality and comprehensive genome-wide protein localisation resource in

**Figure 1.**
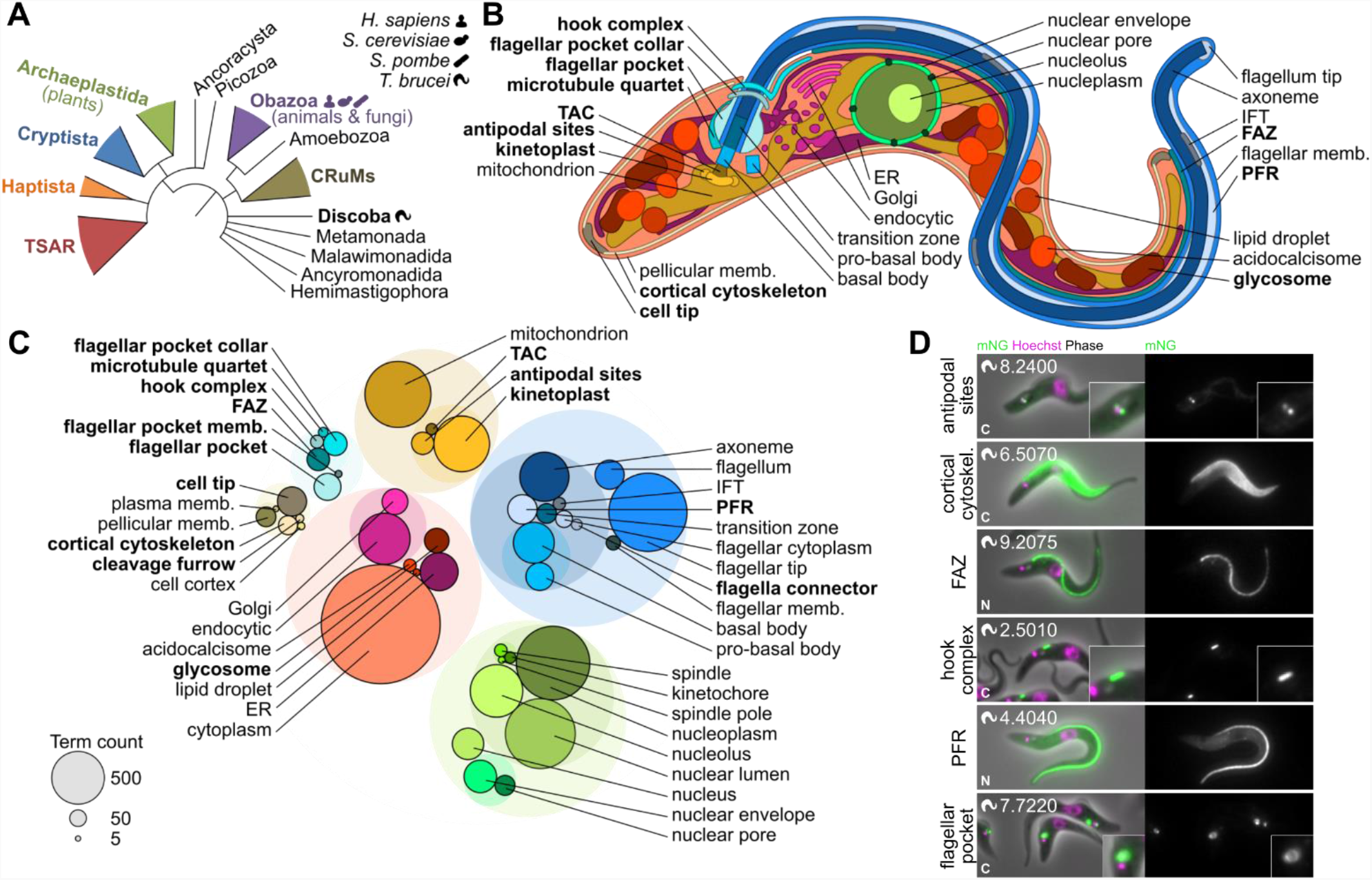
The subcellular protein atlas of *Trypanosoma brucei*. **A**. The position of *T. brucei* in a simplified phylogeny of eukaryotic life, redrawn from Burki et al., 2020. The human and yeast icons are used throughout to indicate when a protein has an ortholog in these species. **B**. The structure of the *T. brucei* cell. Each labelled organelle/structure is distinguishable by light microscopy. Further structures associated with cell division are also distinguishable. Organelles unique to, or with notable elaborations in, the *T. brucei* lineage are shown in bold. **C**. Number of proteins annotated with each annotation term, giving a representation of the relative complexity of each *T. brucei* organelle. Transparent circles represent grouping of annotation terms in an ontology hierarchy. This organelle colour key is used throughout all figures. **D**. Examples of previously uncharacterised proteins localising to different organelles either unique to or with notable elaborations in the *T. brucei* lineage, representative of the quality of microscopy data. *T. brucei* TREU927 gene IDs (minus the Tb927. prefix) are shown in the top left and the terminus of endogenous tagging in the bottom left.

*T. brucei*. We demonstrate the power of this resource for understanding microbial evolution, providing insights to the evolutionary cell biology of eukaryotes and their organelles and show that innovations in organelle complexity around the gain of kinetoplastids parasitic lifestyles are associated with morphogenesis and the cell surface. This powerful single-cell data set also identifies cell cycle stage-specific and organelle subdomain specialisations associated with key parasite functions. The biological knowledge embedded in this data set is a valuable basis for future studies.

## Results

### A high coverage subcellular localisation resource

The *T. brucei* genome^14^ encodes 8,721 proteins, excluding the variant surface glycoprotein (VSG, used for antigenic variation) gene family and identical duplicated genes^15,16^. Using endogenous tagging in the procyclic life cycle stage^17,18^, we generated cell lines and localisation image data for 89% (7,766) of these proteins by N or C terminal tagging and >75% by both N and C terminal tagging (Figure S1A), with ≳250 cells imaged per cell line. Most cell lines had an operationally convincing localisation (see Methods, Figure S1B): 73% of C-terminally and 59% of N-terminally tagged cell lines had greater than background fluorescence signal intensity and/or signal in a position not typical of background fluorescence (Figure S1B). We noted that C or N termini refractory to tagging correlated with known targeting sequences; tagging failures are therefore often biologically informative (Figure S1C,D). We manually annotated protein localisation using a standardised ontology (Figure 1B, Table S1, Table S2) to generate a localisation database (Table S3). Overall, 5,806 (>75% of successfully tagged proteins) had a clear signal (Figure S1A,B), making this a high coverage map and a transformative resource.

This resource maps the protein composition of all organelles (Figure 1C). The most complex were the mitochondrion, nucleus and flagellum, although small organelles also had high complexity (e.g. basal body - 307 proteins). In contrast, the mitotic spindle was simple (30 proteins). Structures adapted or only found in *T. brucei* and related organisms suggestive of parasite specific functions (for examples see Figure 1D) could also be complex, in particular the flagellar pocket (the specialised site of endo- and exocytosis^19^) and the cytoskeleton. *T. brucei* is an early branching eukaryote (Figure 1A), where similarities to its host reflect conserved eukaryotic cell biology^20^ and dissimilarities reflect lineage-specific or parasitism-associated adaptations. For these dissimilar proteins, this resource often provides the first clues to potential function and, importantly, also represent divergent biology which are potential drug targets.

### Parasite-specific and general eukaryotic features

Analysing this unique resource for an early branching eukaryote, revealed *T. brucei* organellar proteins with orthologs across a diverse set of eukaryotes (Table S3, Table S4) indicating the extremely well conserved core organelle machinery (Figure 2A, extended in Figure S2A-E). The nucleus and other membrane-bound organelles (glycosomes/peroxisomes, acidocalcisomes, endoplasmic reticulum (ER) and Golgi apparatus) had a consistently high proportion of conserved proteins (∼25%). The flagellum and mitochondrion had a lower proportion of conserved proteins – likely reflecting innovation in the *T. brucei* lineage. We identified species among these diverse eukaryotes which have lost ancestral features. For example, some (mostly parasitic e.g. *Plasmodium falciparum*) species tend to lack orthologs of *T. brucei* acidocalcisome or lipid droplet proteins pointing to reduced or differing ion homeostasis and variation in lipid metabolism.

**Figure 2.**
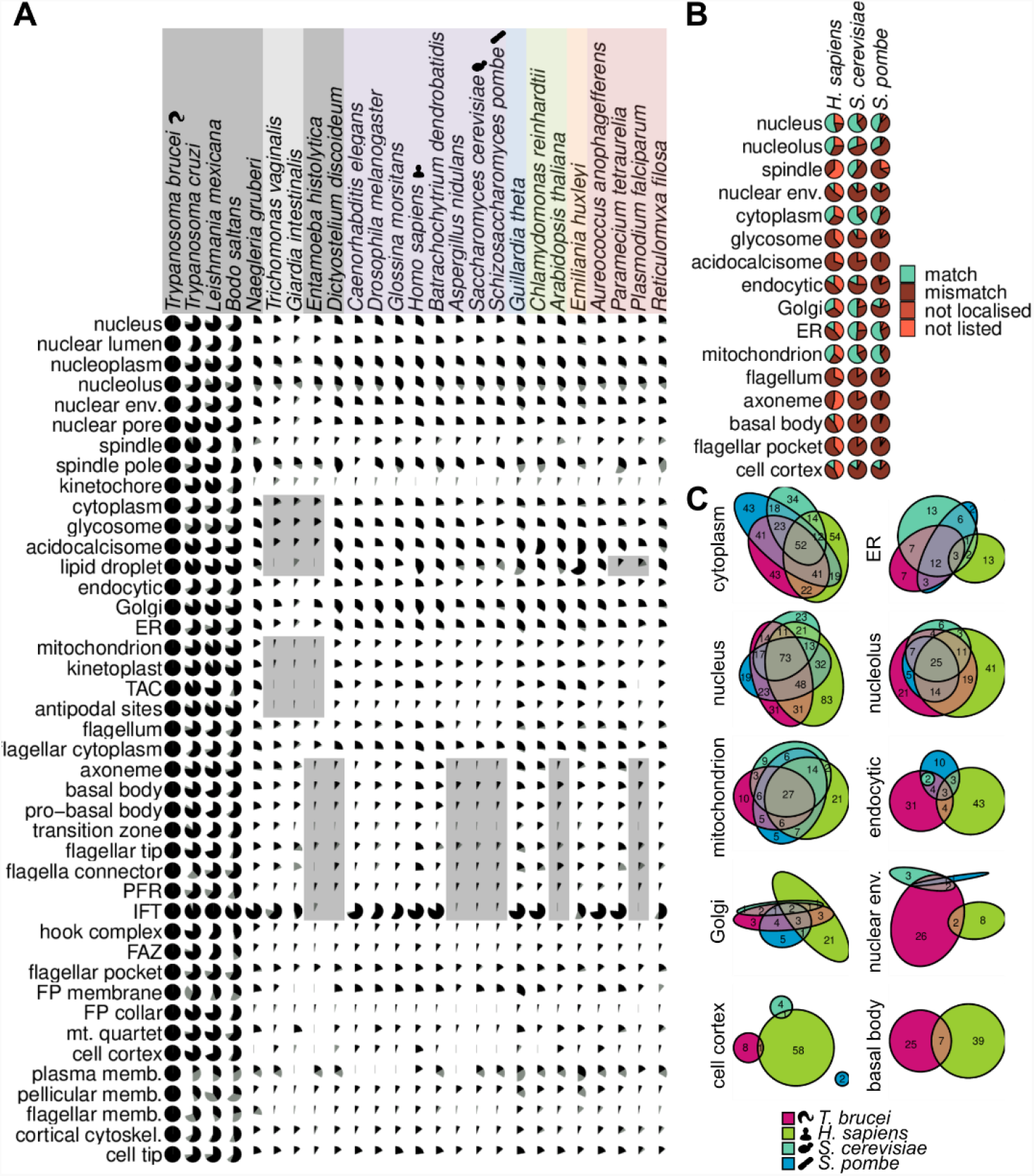
Mapping eukaryote-wide conserved and parasite-specific features. **A**. Presence of orthologs of *T. brucei* proteins, grouped by organelle, across eukaryotic life. Pies represent the proportion of proteins with a reciprocal best BLAST (RBB, black) or not an RBB but at least one orthogroup member (grey) in each species. Extended version in Figure S2. **B**. Conservation of localisation for proteins with an ortholog in humans or yeast (both budding and fission), broken down by organelle, with human/yeast annotations mapped to our most similar *T. brucei* term. Some localisations cannot match (e.g. flagellum) as yeast cells lack the structure and cilia were not annotated in human cells. **C**. Euler diagrams showing the degree of agreement of localisation of the proteins with a single ortholog in human, yeast and *T. brucei* cells or, for the basal body, just humans and *T. brucei*.

Orthologous proteins in different species may act in a different compartment, therefore localisation is also a first indication for whether function is also conserved. Using the comparable human/yeast data^3–5^, we asked whether *T. brucei* proteins had a human or yeast ortholog and whether that protein localised to the same organelle (Table S5, Figure 2B) and, for proteins with a single ortholog in all four species, which orthologs localised to the same organelle in all four species (Figure 2C). Key organelles have conserved proteins with conserved localisation: up to ∼50% localisation conservation in the nucleus and mitochondrion (Figure 2B). However, many organelles do not. As expected, the humans and yeast the cell cortex/surface/cell wall are very different. Furthermore, orthologs of *T. brucei* basal body/centriole, spindle, nuclear envelope and endocytic system proteins tend not to have the equivalent localisation in humans/yeast (Figure 2B,C). This speaks to the vital nature of core nuclear and mitochondrial biochemistry across eukaryotes yet the evolutionary adaptability of the endomembrane system, likely commensurate with the secretory cargo and the specific cell cortex that in *T. brucei* is vital for host-parasite interactions.

Change in localisation presents an opportunity for adaptation, proteins with conserved domains readily assemble into lineage specific structures, such as the extra-axonemal paraflagellar rod^21^ (Figure 2A). Specific protein families tended to have multiple paralogs with a wide range of localisations: calpain-like C2 peptidases, tetratricopeptide and ankyrin repeats (Figure S3A,B).

This is the first genome-wide protein localisation resource mapping the flagellum/cilium – a complex ancestral eukaryotic organelle that is necessary for *T. brucei* morphogenesis and pathogenicity^11,22,23^. The conserved axoneme architecture and evidence from existing high-quality flagellar proteomes^24,25^ point to high evolutionary conservation; therefore, our high-resolution map of flagellar protein localisations will be informative for most flagellated eukaryotes, including pathogens such as *Giardia* and *Trichomonas*. This is also important outside of microbes: in humans, mutations in flagellar/ciliary genes are associated with ciliopathies^26^. There are ∼200 axoneme and basal body *T. brucei* proteins with a human ortholog, many of which are not yet identified as disease-associated (Figure S3C).

### Organelles required for morphogenesis have recent innovations associated with parasitism

This resource allows the first mapping of where and when parasitism-associated adaptations occurred. We determined when protein complexity was gained in each *T. brucei* organelle since divergence from other lineages by identifying divergent species in which an ortholog of each *T. brucei* organellar protein was detected (Figure 3A). Most organelle complexity is either shared with all eukaryotes or arose around the divergence of the class Kinetoplastida. Almost all organelles had a large gain in complexity, with a disproportionately large gain for the mitochondrion, likely associated with evolution of the kinetoplast (the structured mitochondrial DNA). The unusual mitotic spindle and kinetochore also emerged at this time^27,28^. Kinetoplastida includes parasitic and free-living species^29^ so this innovation was not parasitism-associated, but was likely associated with the broader success of this lineage.

**Figure 3.**
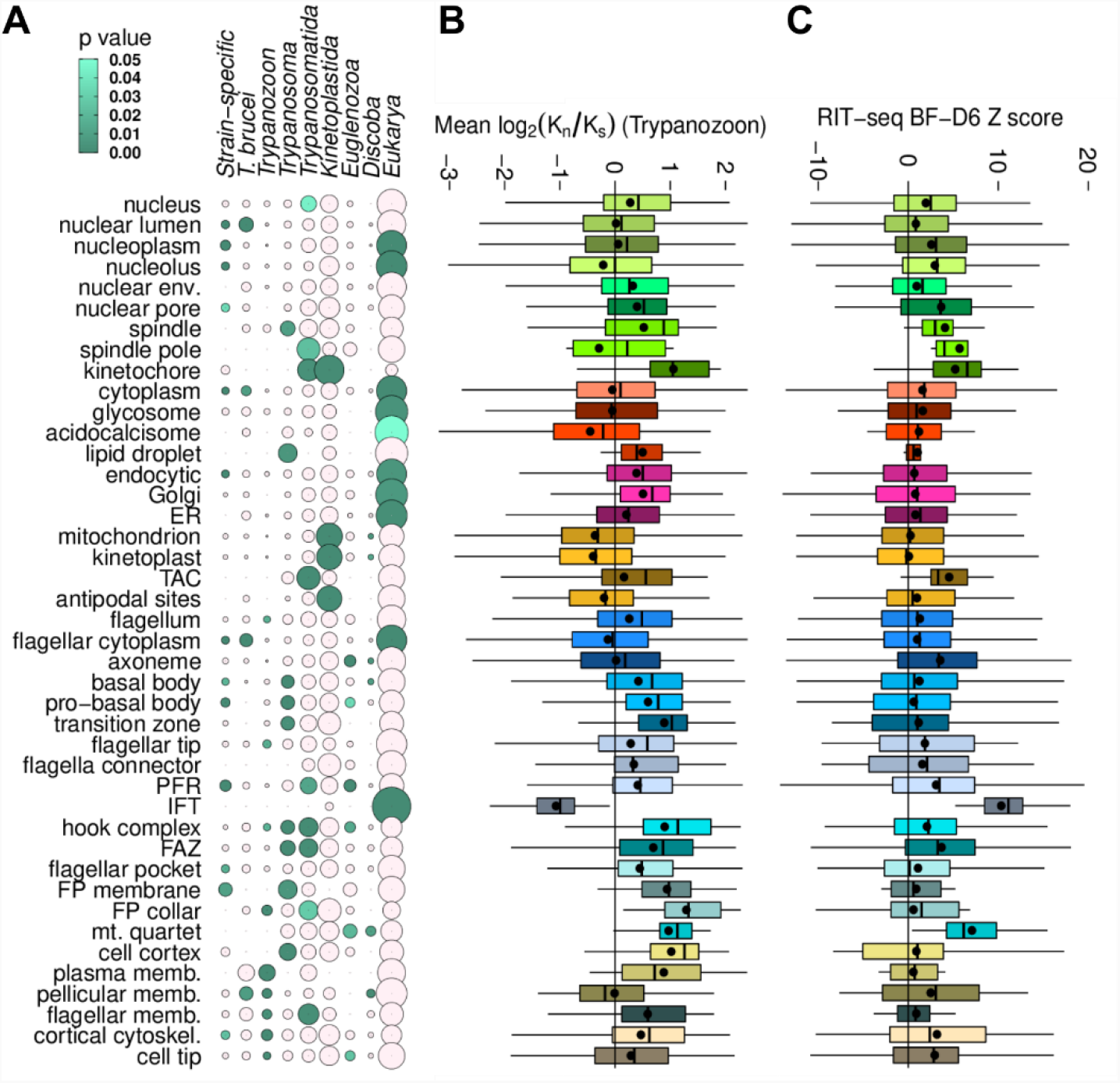
Evolutionary tempos of different organelles reflecting adaptation for parasitism. **A**. Evolutionary distances at which different organelles gained complexity. Circle size represents the proportion of proteins localising to each organelle which have an ortholog (reciprocal best BLAST) in at least one species at that evolutionary distance from *T. brucei* 927 and no detectable ortholog in more distantly related species. Green circles indicate a disproportionately large proportion of organelle complexity gained in or retained from the common ancestor of that evolutionary distance. **B**. Ratio of non-synonymous (K_n_) to synonymous mutations (K_s_) between *T. brucei* 927 and other African trypanosomes (Trypanozoon) for proteins with a single ortholog, broken down by organelle. Among these species amino acid sequence identity is ∼50%. **C**. High throughput fitness phenotype score from ^36^, the result of RNAi knockdown and 6 days in culture as bloodstream forms (BFs), broken down by organelle. Higher values indicate greater fitness cost.

Later adaptations are associated with the evolution of parasitism among Kinetoplastida^30^ and we can now map this to organelles. We also determined the ratio of nonsynonymous and synonymous mutations (*K*_*A*_*/K*_*S*_) per protein among closely related trypanosomes (*Trypanozoon*) (Figure 3B). Mean organelle *K*_*A*_*/K*_*S*_ (> 1 indicating selection for new traits) correlated with recent gain in organelle complexity around the evolution of parasitism (Trypanosomatida) or the evolution of dixenous parasitism (*Trypanosoma*)^29^ (Figure 3A *cf*. Figure 3B). The recently adapted organelles are the cytoskeleton (except the axoneme), plasma membrane domains and the mitotic spindle. This provides new support for evidence linking cytoskeleton mediated morphogenesis to parasitism^19,31–35^.

Mapping RNAi mutant fitness data from the genome-wide *T. brucei in vitro* screen^36^ to organelles (Figure 3C) identified the most vital organelles with least redundancy. Both ancient and more recently evolved structures are vital: the mitotic spindle and kinetochores (more recent), the tripartite attachment complex (TAC, kinetoplast-associated, more recent), IFT (ancient) and the microtubule quartet (MtQ, more recent). These structures are linked with the three sub-cycles which underlie the *T. brucei* cell cycle^13^: nuclear DNA replication and segregation (spindle, kinetochore), kinetoplast DNA replication and segregation (TAC), and the flagellum-dependent cytoskeletal growth and division (IFT, MtQ). Proteins in the spindle, kinetochore, MtQ and TAC are therefore high priority drug targets, exemplified by the recent success of a kinetochore kinase inhibitor^37^. These are on average the most vital organelles; however, parasitism-associated adaptations are likely not limited to these organelles. For example, the cell surface membrane domains also show extensive recent adaptation (Figure 3C).

### Candidate regulators of cell cycle-dependent morphogenesis

The regulation of growth, maturation and size of organelles is a major cell biology question^38^, and understanding these processes will define factors necessary for parasite replication and hence potential targets for intervention. The precisely organised *T. brucei* cell (Figure 1B) and cell cycle means each stage is identifiable from morphology^39–41^. Our resource therefore allows genome-wide molecular-level analysis of organellar growth and duplication and any protein dynamics within an organelle. We found some known^25,42,43^ and many novel proteins which localise specifically to either the newly forming/growing (Figure 4A) or the old/mature (Figure 4B) copy of almost all cytoskeletal structures. This indicates that a different proteomic composition when growing versus mature is a fundamental property of cytoskeletal organelles. These proteins may enable a range of processes, including templated assembly of daughter organelles, enabling/promoting organelle growth, preventing growth, stabilising an assembled structure, or conferring new functions once mature.

**Figure 4.**
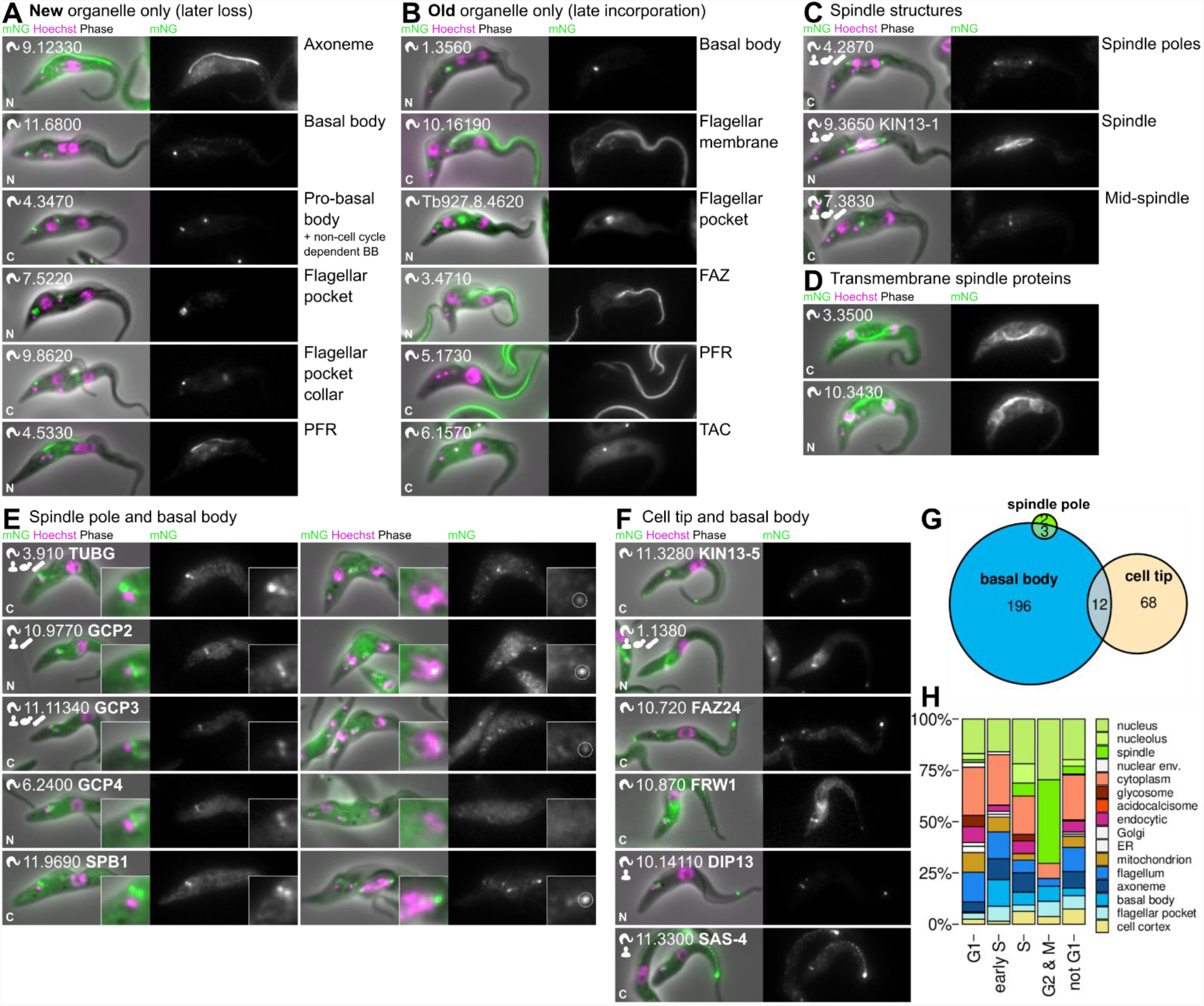
Cell cycle-dependent organelle composition identifies potential morphogenesis regulators. **A**. Six examples of proteins which found more strongly in the new copy of an organelle, showing cells towards the end of the cell cycle (2K1N and 2K2N). **B**. Six examples of proteins which found more strongly in the old copy of organelles, showing cells towards the end of the cell cycle. **C**. Examples of proteins localising to different spindle structures. **D**. Both proteins localising to the spindle with predicted transmembrane domains. **E** Localisation of the proteins which localise to both the spindle pole (spindle nucleating) and basal body/pro-basal body (axoneme nucleating). GCP4 was not detectable at the spindle pole but was included for completeness of the gamma tubulin ring complex. **F**. Localisation of the 6 proteins which localise to both the basal body/pro-basal body (axoneme nucleating) and the cell anterior tip (hypothetically cortical cytoskeleton nucleating) and not to other structures. **G**. Euler diagram of proteins which localise to the axoneme, spindle and cortical cytoskeleton nucleating structures. **H**. Localisation of proteins identified as upregulated at the protein level in particular cell cycle stages by Crozier et al., 2018.

The flagellum and associated structures are critical for *T. brucei* cell morphogenesis. Proteins specific to the new flagellum axoneme and the distal region of the new flagellum attachment zone (FAZ) were particularly numerous. We confirmed several known^44–47^ and identified many novel proteins, including six axoneme proteins. The distal FAZ is the localisation of key cell cycle regulators^44–46^, and the novel components are likely important. Notably, we also identified proteins specific to the flagellar or flagellar pocket membrane associated with either the old or the newly formed flagellum (Figure 4A,B). Protein sorting to plasma membrane domains is therefore sensitive to the maturity of the associated cytoskeleton. This may confer functional differences to cells inheriting the old or new flagellum, important as *T. brucei* life cycle stage transitions are often associated with specialised ‘differentiation divisions’^48,49^.

Mitosis, cytokinesis and mitochondrial inheritance, effected through attachment of the kinetoplast to the basal body and flagellum *via* the TAC, depends on microtubules^13^. Multiple distributed microtubule organising centres (MTOCs) are presumably important for the regulation of these division processes. We identified 318 proteins at the basal body, the MTOC for the flagellum, including 12 of the 15 well-conserved core basal body proteins^50^. The cohort also included 10 kinases and 3 phosphatases, one of which (Tb927.3.690) has previously been identified as important for division^51^. The basal body complexity perhaps reflects its potential as a master regulator of the cell cycle^52^.

*T. brucei* has an unusual chromosomal organisation with many mini and intermediate chromosomes in addition to the 11 megabase chromosomes. These additional chromosomes encode aspects of the VSG gene library on which *T. brucei* antigenic variation is based; however, little is known about their segregation. The nucleus undergoes closed mitosis and, concordantly, the basal body is not associated with the spindle MTOC. Few proteins localised to the spindle pole MTOCs. We identified 14 novel spindle-associated proteins^53^ of which one (Tb927.4.2870, MAP7-like) localised to the poles (Figure 4C). The few other spindle pole proteins are known: the γ-tubulin ring complex (γTuRC), MLP2 and SPB ^53–56^, constraining possible mechanisms for intermediate/mini-chromosome segregation. Two proteins (Tb927.10.3430, Tb927.3.3500) had transmembrane domains (Figure 4D) and one (Tb927.11.5110) also localised to the nuclear envelope, possibly indicating roles in closed mitosis.

Different trypanosomatids have distinctive morphologies, likely sculpted by the interaction with the host and vector^33^, defined by the sub-pellicular microtubule array which is a parallel one-layer thick corset of microtubules under the plasma membrane^57,58^. How the microtubules within this array are nucleated remains unknown: perhaps via dispersed nucleation within the array or via a major anterior MTOC where a subset of microtubules ends are located, then movement into the array to add new microtubules and maintain spacing as the cell widens. As previously described^55^, we did not detect γTuRC proteins in the array (Figure 4E). Interestingly, at the sensitivity we achieved, we identified no proteins shared among all MTOCs. Several proteins including SAS-4 (Tb927.11.3300) localised to the basal body and the sub-pellicular array (Figure 4F), but we identified none which also localised to the spindle poles (Figure 4C-G). We identified many sub-pellicular array-associated proteins, these may contribute to nucleation or organisation. ∼60 proteins localised to most of the array and many more localised to one of the ends the array at the cell tips (Figure S4). As the tip of the new growing FAZ becomes the site of furrow ingression^57^ and this new sub-pellicular array anterior tip is likely a key MTOC for sub-pellicular microtubules.

Like other eukaryotes, the *T. brucei* cell cycle is regulated by cyclins (CYC) and cyclin-related (CRK), aurora (AUK) and mitogen activated (MAPK) kinases^13^. However, it is incompletely known how these proteins control division of the *T. brucei* cell architecture. Previous meta-analyses, for example of the spindle assembly checkpoint^59^, were limited to ortholog presence or absence, while this localisation implicates some proteins in regulation of division of specific organelles: MAPK4 (Tb927.6.1780) localised to the basal body, MAD2 (Tb927.3.1750) to the microtubule quartet, MAPK6 (Tb927.10.5140) to the cortical cytoskeleton and posterior cell tip and CRK1 (Tb927.10.7070) to the mitochondrion. CRK1 location maybe particularly important as it is unknown how the division of the mitochondrion and kinetoplast are coordinated. Proteins with cell cycle-dependent abundance^60^ tended to localise to organelles dividing at the corresponding cell cycle stage (Figure 4H). Some proteins had localisations which changed through the cell cycle; however, we only re-identified proteins with known dynamics: AUK1 (Tb927.11.8220), CYC6 (Tb927.11.16720), and AUK3 (Tb927.9.1670)^61–63^. These remain the most interesting candidates for master cell cycle regulators.

### Novel organelle subdomains: key parasite functions and life cycle transitions

Specialised organelle subdomains often carry out specific functions and through our resource we discovered that most *T. brucei* organelles have such subdomains. Presence of subdomains points to dynamic processes assembling or maintaining them, as a uniform protein distribution typically reflects random processes like free diffusion.

*T. brucei* flagellar-driven motility is critical for virulence in the mammalian host and development in the tsetse fly^64,65^. The flagellum and other cytoskeleton-based structures have important functions associated with specific subdomains^25,46,47,66–68^ and we identified numerous proteins associated with axoneme, cortical cytoskeleton and FAZ subdomains (Figure S4). In particularly, we showed that the flagellum tip was a complex structure (Figure S4A). 60 proteins localised to either the axoneme tip or the flagellar membrane tip. The latter is of interest for environmental sensation as *T. brucei* swims with the flagellum leading. We re-identified known proteins^69,70^, and identified several novel proteins plausibly involved in signalling: Casein kinase (CK1, Tb927.3.1630), a META domain-containing protein (Tb927.5.2230), whose ortholog in *Leishmania* is a virulence factor^71^, and a sodium/hydrogen antiporter (Tb927.11.840).

*T. brucei* has a number of life cycle developmental forms with characteristic morphologies and substantial remodelling of the cortical cytoskeleton is required for their generation. We identified 10 subdomains of the cortical cytoskeleton (Figure S4D,E), greatly extending the previous discovery of a posterior-ventral domain marked by PAVE1 (Tb927.8.2030)^67,68^, with particular complexity seen in the anterior and posterior tip subdomains. Like the flagellum tip, the cortical cytoskeleton posterior tip maintained its protein cohort over the cell cycle. These proteins tended to have a point-like or a cone-shaped signal (see below). As the posterior tip included known end-binding/tip-tracking EB1 (Tb927.9.2760)^72^ and XMAP215 (Tb927.6.3090)^73,74^ and many kinesins, their dynamics likely maintain its structure. The anterior tip was comparably complex, with kinesins, known vital kinetoplastid-specific TOEFAZ/CIF components and proteins also specific to the distal FAZ^44–46^ (Figure S4E). Together, this indicates that the cortical cytoskeletal is far more complex and dynamic than previously appreciated.

*T. brucei* is an exclusively extracellular parasite and throughout its life cycle, the parasite surface is predominantly covered in GPI-anchored membrane proteins. Beyond the well characterised ER exit site (ERES) and nuclear envelope, further ER compartmentalisations were detected (Figure 5A-E). These domains likely represent functional specialisations as, for example, the GPI transamidase complex necessary for the GPI anchoring of the dominant surface protein localised to a distinctive cisternal-like subdomain (Figure 5B *cf*. Figure 5C). A subset of chaperone proteins, DnaJ46 (Tb927.3.1430) and BiP (Tb927.11.7460), had a similar localisation. Our observations illustrate the level of complexity within the ER, bringing into focus the challenge of understanding how a cell secures or actively maintains enzyme position in subdomains as they process a high flux of substrate.

**Figure 5.**
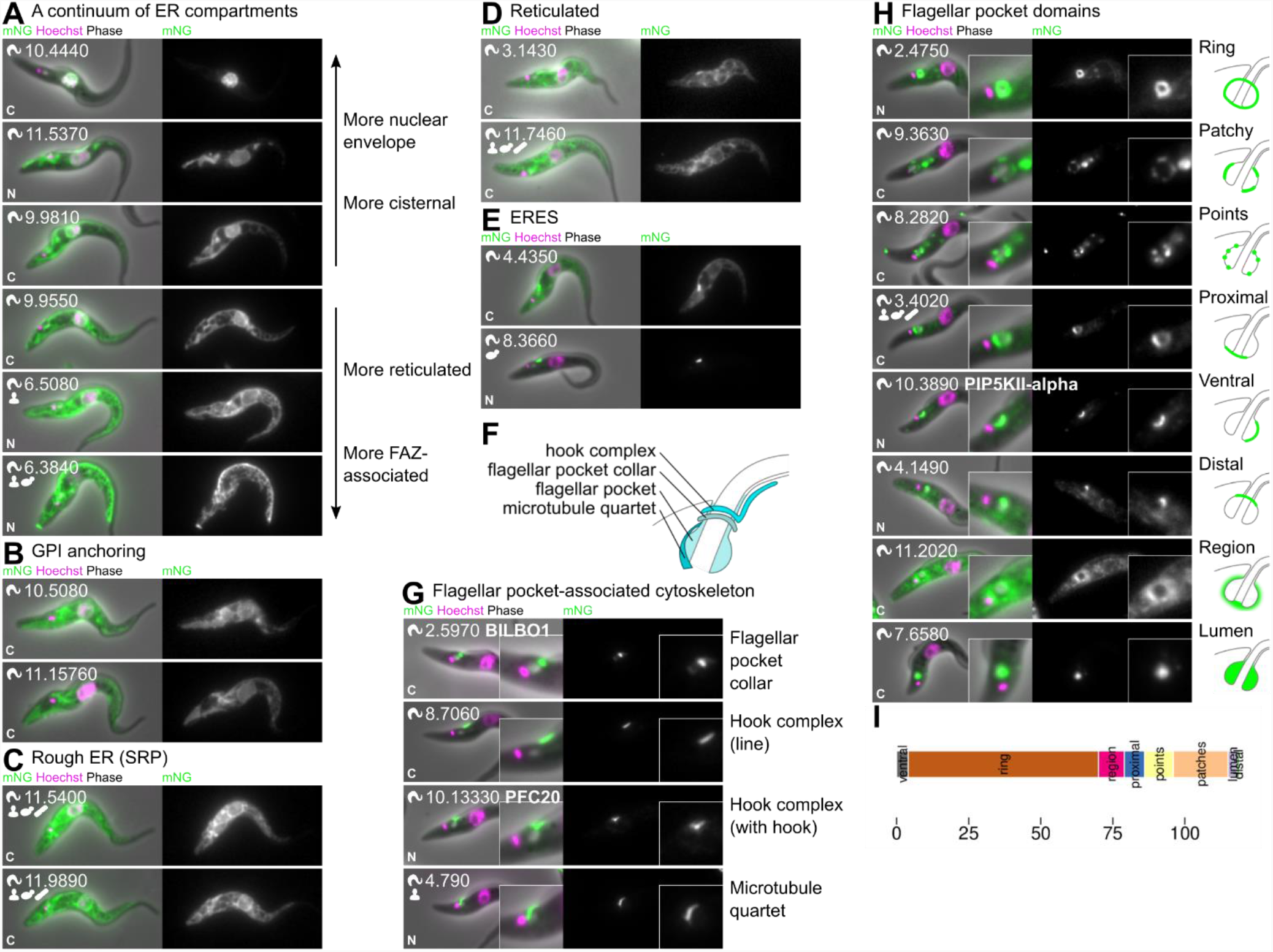
Specialised subdomains of the endomembrane and endo- and exocytic systems. **A**. Examples of proteins localising to different regions of the endoplasmic reticulum, which broadly exist as a continuum from more cisternal to more reticulated. **B**. Enzymes for GPI anchoring tend to have a moderately cisternal localisation. **C**. Signal recognition particle (SRP) proteins tend to have a moderately reticulated localisation corresponding to rough ER. **D**. Examples of proteins with a particularly reticulated localisation. **E**. A large subset of proteins also localise strongly to the ERES in addition to the ER. **F**. Cartoon of the cytoskeleton structures associated with flagellar pocket. **G**. Examples of proteins localising to cytoskeleton structures associated with the flagellar pocket. **H**. Examples of proteins localising to different flagellar pocket domains and flagellar pocket-associated domains. **I**. Number of proteins localising to each flagellar pocket sub-compartment.

High flux is also necessary to maintain the cell surface, with rapid trafficking of material onto and off the host-exposed cell membrane. Specialisation of membrane domains extends to the site of endo- and exocytosis and complexity in these systems is common^75^. We found that the flagellar pocket, supported by a specialised, highly complex cytoskeleton (Figure 5F,G), has 8 characteristic subdomains (Figure 5H,I), for example a lipid kinase (Tb927.10.3890) localised to the ventral flagellar pocket. These pocket subdomains potentially support the high endo- and exocytic flux through this small membrane domain^76^. We identified two apparent lumenal flagellar pocket proteins (Tb927.7.6580, Tb927.7.6590), showing that *T. brucei* maintains a small, localised extracellular environment.

Gene expression control in *T. brucei* has a number of unusual features, with co-transcribed gene arrays with subsequent message processing producing mature mRNAs by addition of a splice leader to the 5’ end, and the use of RNA pol I to transcribe the mRNA for the surface coat proteins, procyclin and VSG. We identified cohorts of proteins localising to characteristic patterns of 1, 2, or multiple points within the nucleoplasm, in addition to nucleolus-associated points, (Figure S5A-C). Based on previously characterised proteins, these patterns are likely associated with structures such as Pol II factories for spliced leader RNA transcription^77^, telomeres^78,79^, and fibrillarin-like (potential Cajal bodies)^80^. *T. brucei* RNA polymerase I (Pol I) subunits (RPA proteins) tended to localise at the nucleolar periphery and basal Pol I transcription factors (CITFA proteins) localised to points around the periphery, leading to a prediction that proteins with a similar localisation will have a Pol I associated function. Overall, the organisation of the trypanosome nucleolus differs from that of metazoa^81^.

In contrast to the ER, we found the mitochondrion to be uniform in composition. However, the kinetoplast region appeared complex (Figure S5E,F) with known and novel proteins which localised to the antipodal sites^82,83^ (associated with DNA replication), localised to the TAC^84–86^, and co-localised with kinetoplast DNA^87,88^. We also identified novel kinetoplast-associated foci, although their potential function is cryptic.

### Cell posterior is a site of complexity and protein moonlighting

Our genome-wide *T. brucei* protein localisations built a picture of a eukaryote with all the normal organelles, but distinctive specialisations, yet the posterior cell tip stood out as a defined structure of unexpectedly high complexity (Figure 6). Many proteins localised specifically to the posterior tip but it was also a common second site for proteins localising to another organelle or structure. This may be a ‘moonlighting’ localisation in addition to the expected or known site of function of a protein.

**Figure 6.**
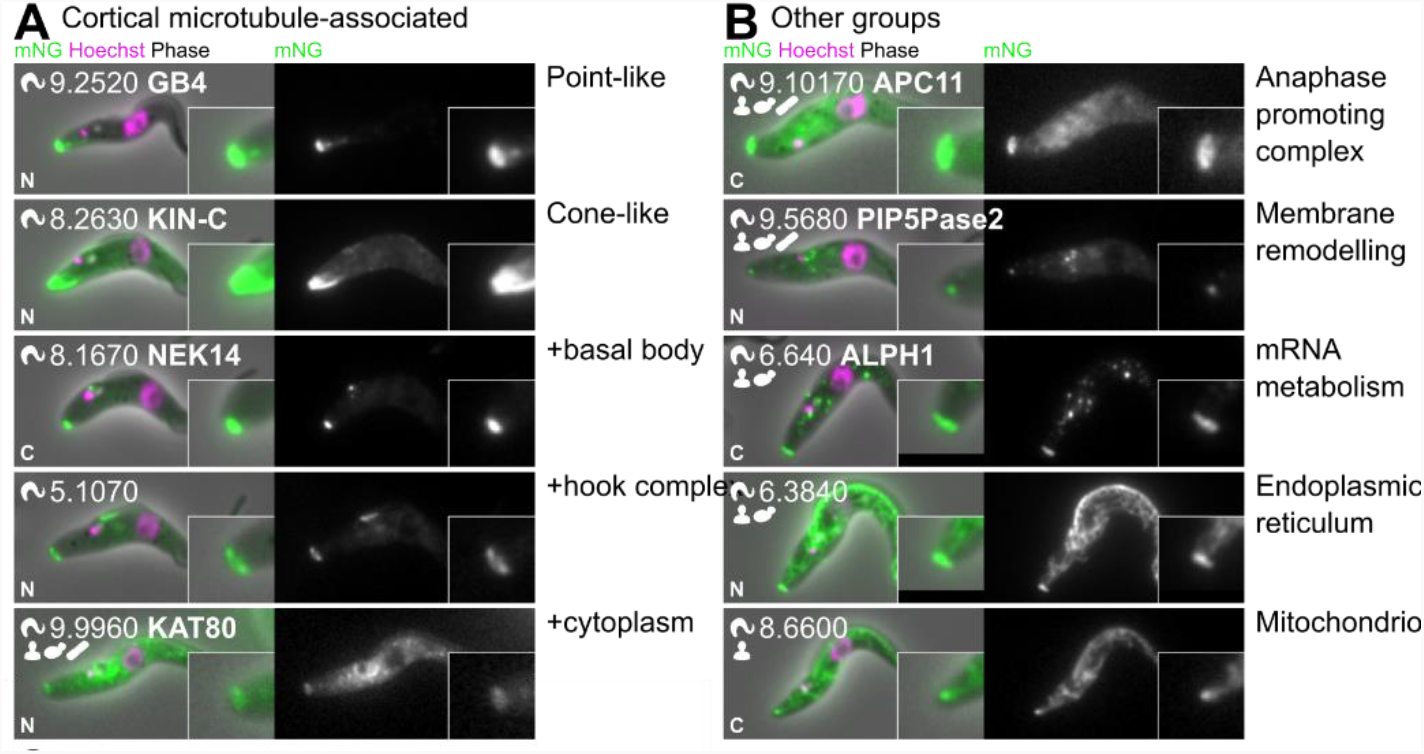
Proteins involved in diverse processes also localise to posterior cell tip. **A**. Many proteins which localise to the posterior tip are plausibly microtubule associated and a subset also localise to other microtubule-containing structures. **B**. Proteins with diverse predicted functions strongly localise to the cell posterior tip, with some also localising to other structures in the cell.

The posterior tip therefore has linked or related functions with several organelles. Firstly, many microtubule associated proteins localised to various cytoskeletal structures and the posterior tip (Figure 6A). Secondly, proteins associated with endo- or exocytosis (including clathrin), often localised to the flagellar pocket with an additional focus at the cell tip, perhaps for roles in membrane remodelling or an alternative site of import/export. Thirdly, proteins which localised to the ER or mitochondrion (including DLP1) but with a focus at the cell tip, indicating a site of ER/mitochondrion-posterior interaction or membrane management (Figure 6B). Fourthly, proteins likely involved in RNA catabolism, including two known (XRNA^89^ and ALPH1^90^) and seven novel, mostly found in RNA granules but with a focus at the cell tip. Finally, 9 out of 10 anaphase promoting complex (APC) proteins^91^ and the APC-interacting kinetochore protein KKT10 (Tb927.11.12410)^92^ localised to the posterior cell tip, in addition to the nucleus during mitosis (Figure 6B). The functional importance of this is unclear: is the cell posterior just a common site for sequestration? However, the proteins present at the posterior suggest cell cycle-dependent membrane management for abscission of the plasma, ER and mitochondrial membranes.

## Discussion

Protein localisation, and timing of localisation, is central to unicellular parasite cell function, defining the site of action of a protein, the substrates or interaction partners available and when they may act/interact. Determining localisation from microscopy is particularly powerful, being high content single cell data. Our mapping of the *T. brucei* cell comprises not only 7,766 protein localisations but also ∼5,000,000 individual cell images, capturing cell cycle and organelle dynamics, available for ongoing analyses.

*T. brucei* is a highly structured single cell organism, illustrative of the enormous diversity among unicellular eukaryotes and analogous to other unicellular parasites like *Giardia, Trichomonas* and *Plasmodium*. Our localisation data indicates extensive placement of specific biochemistry in specific sites or organelle subdomains, for example the complexity of the flagellar pocket membrane or the multiple specialisations of the cell posterior. Nonetheless, even in this early-branching lineage, much eukaryotic biology is conserved and informs our understanding of eukaryotic evolution. As protein structure predictions^93,94^ improve our ability to detect extremely divergent orthologs, *T. brucei* will become even more informative for understanding both fundamental biology and pathogenic specialisations.

Some specialisations reflect adaptations for parasitism and often represent streamlining or examples of “extreme biology” where normal structures become highly elaborated. Our data show extensive recent morphology-associated adaptations in the cytoskeleton and its dynamics, especially associated with the specialised endo-/exocytic organelles. This defines the molecular machinery responsible for the morphological specialisations, which are a defining feature of the trypanosomatid parasites. Our data hint at new adaptation themes. Even in the procyclic form, not directly exposed to an adaptive immune system, the exposed cell surface is simple compared to the flagellar pocket - perhaps some environmental sensation occurs in the flagellar pocket and endocytic system? The spindle has undergone many recent changes - is this associated with the evolution of the library of VSG-containing minichromosomes? Our localisation database is a key resource for further work to understand adaptation for parasitism.

Protein localisation data has a key intrinsic limitation that the protein must be expressed to be localised and our data reflects the *T. brucei* procyclic form expression program. This caveat applies to life cycle-specific expression in other unicellular organisms, such as spores and meiotic stages in yeast and the multitude of highly adapted life cycle stages in apicomplexa. This is therefore a generic issue for protein localisation databases in all systems. For *T. brucei*, procyclic non-expression may be evidence that the protein has stage-specific expression. Use of this information identificatied the first bloodstream form-specific transcription activator for monoallelic antigen expression, ESB1^95^.

Localisation may also be influenced by the position of the tag. Tagging may disrupt cryptic post-translational modifications, targeting sequences, or only localise part of the protein if proteolytically cleaved. Also, spatial positions are informative: an example being the large TAC protein p197 where the visibly different but adjacent localisations by N (TAC) and C (basal body and pro-basal body) terminal tagging likely reflect its physical size and orientation.

The value of genome-wide protein localisation impacts all organelles, particularly in a highly structured cell like *T. brucei*. Our database is important for interpreting how cellular complexity is encoded by a genome and is modulated in space and time. *T. brucei* is now the fourth eukaryote, the first eukaryotic pathogen, and the first flagellate for which a genome-wide protein localisation resource has been constructed. The http://tryptag.org database contains information on all organelles and is an important searchable resource and the images and associated annotations have been integrated onto VEUPathDB^16^. The addition of an early-branching eukaryote to the other three organisms is of particular value, enhancing our view of general eukaryotic and parasite evolution.

## Methods

### Cell lines, culture and genetic modification

*Trypanosoma brucei brucei* TREU927 was selected for our protein localisation database as it is the original genome strain ^14^ with a high quality reference genome supported by community annotation ^15,16^. We used the SmOxP9 line, which also expresses T7 polymerase and Tet repressor ^96^. The procyclic form life cycle stage was used as it is readily grown in culture, grows to high densities and is more amenable to high throughput tranfection efficiencies. Procyclic forms were grown in SDM79 ^97^.

Tagging constructs were generated using a long primer PCR from a standard plasmid (pPOT v4) encoding a drug selectable marker (BSR) and a fluorescent protein with GS linkers (mNG), as we previously described ^17^. The 5′ primer ends are 80 bases of homology allowing homologous recombination into the target locus, when introduced by electroporation ^17^. Long primer PCR, electroporation and drug selection in multi-well plates were carried out as we previously described ^18^, with the overall plate workflow following our project announcement ^98^. Following electroporation in 96 well plates, the transfectants were then transferred to 4 24 well plates for drug selection of stable transfectants in 2 ml culture medium with 20 µg/ml blasticidin. Once transfectants had grown to ∼4×10^7^ cells/ml they were subcultured once in 24 plates to approximately 1×10^6^ cells/ml, followed by 24 h growth was used to give a healthy population for microscopy

During high throughput tagging, success rates (percent success in generating a construct by PCR, selecting a drug resistant cell line, and observing a convincing subcellular localisation) were monitored (Figure S1A-C), as was agreement with known localisations and existing proteomic data (Figure S1E). Failures were repeated at least once, and as repeats had a comparable success rate to first attempts, appeared largely stochastic (Figure S1A). Some genes were truly refractory, including those with known problematic gene models, for example Tb927.11.1090 C-terminal tagging consistently failed. This gene model is actually the N terminus of GM6, an extremely large and repetitive gene, with C terminal GM6 found in the downstream gene model Tb927.11.1100 ^99^). Such gene model issues were not corrected here.

Microscopy was carried out on live cells. As they are motile, the cells were washed three times with vPBS which allows them to adhere to glass and 500 ng/ml Hoechst 33342 was included in the first wash to stain the DNA ^100^. Micrographs were captured on a Leica DM5500 B epifluorescence microscope with a 100W mercury arc lamp and either a 100× or 63× NA1.4 oil immersion objective (the vast majority at 63×) and an Andor Neo 5.5 sCMOS camera running in 16-bit high well capacity mode, using Micromanager ^101^. mNG fluorescence was captured using the L5 filter cube, excitation 480/40 nm, dichroic 505 nm, emission 527/30 nm. A standard exposure time of 2000 ms was used, unless the mNG signal was particularly bright. Typically, 4 to 5 fields of view (aiming for 200 or more cells) were captured, with more when a cell line was identified as having a rare (i.e. cell cycle dependent, like spindle) signal.

Before analysis, all images were subjected to the same corrections: First, camera amplifier offset per pixel column in the upper and lower halves of the images (measured from no illumination images) were subtracted. Second, the median of all images captured on that day (typically >200) was taken as background signal and used for flat field correction. Then, finally, scaling of 100× images to 63× equivalent, and normalisation the green channel signal intensity from exposure time and any magnification scaling (2.51×).

Microscope images were manually checked for quality (healthy cell appearance, cell number, appropriate exposure time, focus, etc.) and the tagging flagged for repeating if there were significant quality concerns.

### Gene selection for tagging

Tagging was based on version 5.1 of the genome sequence ^14^. The initial target set based on TbruceiTREU927 genome annotations from TriTrypDB release 5.0 (June 2013). This comprised all genes which mapped to one of the 11 megabase chromosomes (Tb927_01_v5.1 to Tb927_11_v5.1), excluding the chromosome 11 right hand fork (Tb927_11_RH_fork_v5.1) and excluding genes annotated as a variant surface glycoprotein (VSG) or VSG expression site associated genes (ESAGs). The latter are only expressed in the bloodstream form, where VSGs are used for antigenic variation.

New gene models added by TriTrypDB to the megabase chromosomes over the course of the project up to TriTrypDB release 45 (August 2019) were also tagged, as were previously identified transcribed small ORFs ^102^ (which could be unambiguously mapped to the genome) and a small number of manually selected genes not mapped to megabase chromosomes. We attempted tagging of 479 genes whose gene models were removed as unlikely as of TriTrypDB release 45. Strain-specific proteins tended to localise to the nuclear lumen, cytoplasm and flagellar cytoplasm – this localisation can arise spuriously (Figure S1D) and we suspect that these are dominated by incorrect gene models.

Integration of the transfected construct occurs using the endogenous homologous modification machinery, specificity is therefore conferred by the uniqueness of the homology arms. Generally, only one N and one C terminal tagging attempt was made for a set of genes which could not be uniquely targeted. This mostly affected genes which are found tandemly duplicated or in an array.

All genes were tagged at the C terminus, irrespective of whether they had a predicted targeting sequence (e.g. glycosomes have a known C terminal targeting tripeptide^103^). Genes not predicted to have an N terminal signal peptide (SignalP *p*<0.5, as indicated by TriTrypDB at the time of primer design) were tagged at the N terminus. Note that this is not a sensitive predictor in *T. brucei*.

### Annotation of protein localisation

Each cell line was manually annotated by a group of at least 3 experts using an ontology of 45 annotation terms for different organelles/cell structures. This extended our previous description of the characteristic appearance of over 30 different organelles and structures ^74^. This hierarchical ontology has specific (e.g. ‘nucleoplasm’ or ‘axoneme’) and more general terms localisations (e.g. ‘nucleus’ or ‘flagellum’). The more general term was used if a localisation was ambiguous. If localising to multiple organelles then all relevant terms were used (i.e. additive annotation) (Table S1).

We used a further ontology to describe any additional structure within each organelle (e.g. ‘patchy’, ‘weak’ or ‘points’) ^74^. In some cases, these were used for lower confidence annotations (e.g. Nucleus [points] rather than nuclear pores). The ‘weak’ modifier was reserved for localisations with signal comparable to background autofluorescence (Table S2).

Manual annotation weak signals were supplemented by automated mNG fluorescence signal intensity, measured from all cells from all images of all cell lines. For reference, 4 independent samples of the parental cell line were grown and prepared for microscopy identically to tagged cell lines. Individual cells were identified, oriented and cropped from the images automatically using intensity thresholding of the phase contrast image after a series of unsharp and background subtraction filters to generate cell masks, as previously described ^104,105^ and mean mNG signal intensity (sensitive to overall signal) and 99^th^ percentile mNG signal intensity (sensitive to small bright structures). Autofluorescence tended to occur in the mitochondrion, cytoplasm and/or endocytic system. Therefore, any mitochondrion, cytoplasm or endocytic system annotation where both mean and 99^th^ percentile green signal intensity were below the parental cell line were automatically given the ‘weak’ modifier. mNG images are displayed mapping black to the median signal outside of cells and mapping white to 4,500 or the maximum pixel value, whichever is higher.

Cell lines were non-clonal, as necessitated by the high throughout. From previously determined transfection efficiency ^17^, we estimate that populations were typically derived from 5 to 20 clones. In some cell lines this leads to heterogeneity, and in these cases organelle annotations were given a modifier of the approximate proportion of the population with the signal.

For all downstream analyses, a protein was listed as a component of an organelle if it was annotated as localising to that organelle/cell structure, any substructure of that organelle, not annotated ‘weak’ and occurring in at least ∼10% of the population. For some analyses, a simplified set of localisations are used. In these cases, proteins were listed as localising to the nearest parent term in the simplified list.

This database of microscopy data and human annotations can be viewed and downloaded at http://tryptag.org with the annotations and an example image viewable and searchable at http://tritrypdb.org.

### Evaluation of localisation reliability

Localisations for a protein from N and C terminal tagging represent independent biological replicates. In most (>70%) cases both termini gave a similar localisation (Figure S1C). In the remainder, one terminus tended to give no detectable signal while the other gave a clear localisation, as noted in the results this often correlated with a known targeting sequence. Only in a small minority (2%) was there a clear discrepancy (Figure S1C). Therefore, no localisations were excluded from genome-wide analysis.

mRNA UTRs, particularly 3′ UTRs are implicated in life cycle stage regulation. We tested the impact of 5′ UTR (by N terminal tagging) and 3′ UTR replacement (by C terminal tagging) using known life cycle stage specific paralogous protein pairs, which showed expression levels which correlated with the expected stage specificity despite UTR replacement (Figure S1F). Some cell lines with undetectable signal may represent specific expression of that protein in a different life cycle stage.

We treated two localisation types with lower confidence, as they tended to occur as a ‘contaminant’ at low frequency in cell lines: No detectable signal (‘faint’ cells, similar to the parental cell line) or cells with a uniform cytoplasm, nuclear lumen and flagellar cytoplasm signal (‘bright’ cells, similar to cells expressing mNG not fused to a protein). To determine their origin, we cloned and sequenced the modified locus in one ‘faint’ and one ‘bright’ contaminant from otherwise successful tagging. Both had a frame shift originating from the plasmid binding region of the primer, in both cases introducing an early stop codon. They are therefore stochastic errors likely arising from primer synthesis errors.

### *Trypanosoma brucei* protein properties

Gene metadata for analysis was derived from TriTrypDB release 47 (April 2020). This includes gene names and descriptions, genome coordinates, predicted gene mRNA, CDS and protein sequences and basic predicted protein properties (molecular weight, isoelectric point, etc.). Predicted functions were derived from predicted PFAM ^106^ and SUPERFAMILY ^107^ protein domains via TriTrypDB.

Pol II protein-coding gene transcription units were mapped manually. Genomic regions over 50 kbp with non-VSGs genes in a consistent orientation were mapped as transcription units, ignoring occasional genes in the opposite orientation in a transcription unit unless there was high plausibility of expression (e.g. rRNA genes or protein-coding genes with a clear localisation). Transcription units comprising >20 protein-coding genes with localisation data were analysed.

Predicted mitochondrial presequence and signal peptide targeting sequences were identified using TargetP-v2.0^108^.

### Orthology and selection pressures

Evidence for evolutionary trends and conservation in specific organisms was derived from predicted protein sequences from 102 genomes, with broad coverage of eukaryotes and comprehensive coverage of Discoba lineages. This was supplemented with 10 transcriptomes from Excavata lineages with poor genomic coverage, from which protein sequences were predicted using TransDecoder v5.5.0 (LongOrfs)^109^ (Table S4).

Orthologous groups were determined Orthofinder v2.3.12 ^110,111^ using default settings using diamond v2.0.5 and FastME 2.1.4. Reciprocal best BLAST (RBB) hits to *T. brucei brucei* TREU927 proteins were identified using NCBI BLAST 2.9.0+ ^112^, reciprocal hits irrespective of forward and reverse search e-value and accepting reciprocal hits which identified a *T. brucei* gene with an identical sequence to the starting gene. For our analyses, a protein in a different species was defined as ‘the’ ortholog of a *T. brucei* gene if it was either the RBB or was the only orthogroup member in that species. We have used current bioinformatic approaches for all protein-protein ortholog analyses but these are limited by the power of such computational comparative approaches.

Gain in complexity at different evolutionary distances was carried out using NCBI Taxonomy species classifications. We scored the proportion of proteins localising to a particular organelle which had an ortholog in at least one species at that evolutionary distance (Table S4) using a hypergeometric test to detect over-enrichment.

To determine the ratio of number of nonsynonymous mutations (*K*_*A*_) to synonymous mutations (*K*_*S*_) RBB protein sequences were aligned using Clustal Omega ^113^. The corresponding CDSs were mapped to the protein sequence alignment and scored for identical codons (no mutation), synonymous mutation, nonsymonymous mutation or indel mutation (alignment gap). *K*_*A*_*/K*_*S*_ was calculated per codon treating gaps as nonsynonymous mutations. *K*_*A*_*/K*_*S*_ was calculated without any codon bias correction for *T. brucei brucei* TREU927 against each *Trypanozoon* (African trypanosome) species for each reciprocal best BLAST ortholog, and averaged.

### Human and yeast proteins

Human/yeast protein localisations were obtained from the respective project websites. For C terminal whole-genome yeast tagging projects ^3,114^, https://yeastgfp.yeastgenome.org/allOrfData.txt for *S. cerevisiae* and a custom web scraper from https://www2.riken.jp/SPD/01/01A01.html for *S. pombe* (both accessed December 2020). For the human antibody-based Human Protein Atlas/Cell Atlas project ^5^, https://www.proteinatlas.org/download/subcellular_location.tsv (accessed September 2020). Annotation terms were remapped to the most similar *T. brucei* structure for comparison (Table S5).

Evidence for involvement in human genetic disease was determined from OMIM ^115,116^, accessed November 2018. Entries per gene were mapped to Ensembl gene IDs (using the OMIM-provided mapping) and Ensembl gene IDs were mapped to Uniprot protein IDs (using the Uniprot protein mapping). Proteins were taken as involved in disease if the parent gene was annotated as associated or statistically linked with disease, involved with a known molecular mechanism or involved along with multiple genes.

## Acknowledgements

We would like to thank the Wellcome Trust for funding through Investigator Awards to KG and MC [104627/Z/14/Z, 217138/Z/19/Z] a Biomedical Resource Grant to KG [108445/Z/15/Z], a Sir Henry Wellcome and a Sir Henry Dale Fellowship to RW [211075/Z/18/Z, 103261/Z/13/Z], and a Biomedical Resource Grant supporting VEuPathDB [218288/Z/19/Z]. We would also like to thank the TrypTag scientific advisory group for their support and advice.

---

**Figure S1.**
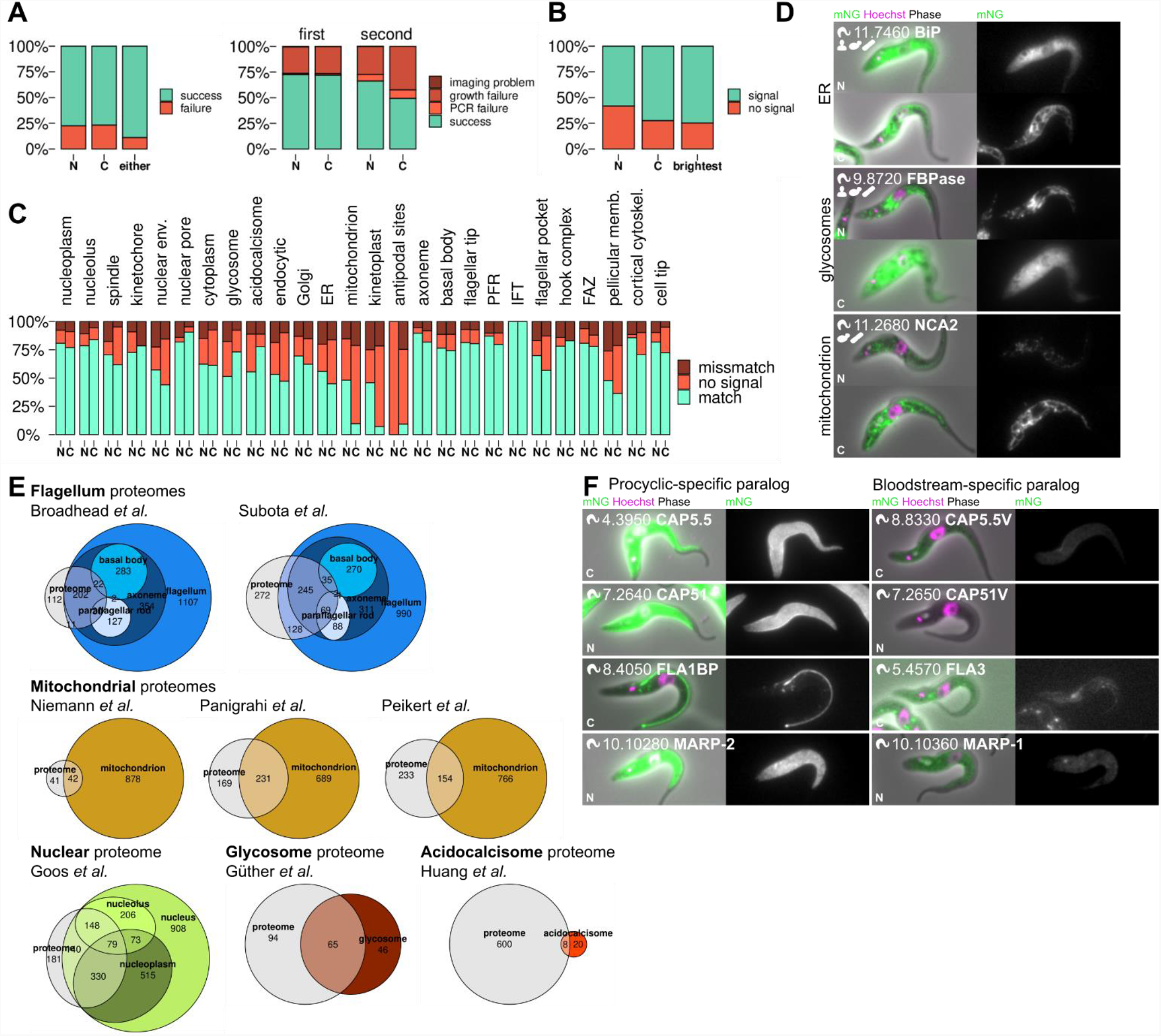
Localisation success rates and reliability. **A**. Left, the proportion of proteins for which a tagged cell line and microscopy data was successfully generated from N terminal tagging, C terminal tagging and from at least one (either) terminus. Right, the success rates from the first attempt and a repeat attempt at endogenous tagging, leading to the overall success rate on the left. **B**. The proportion of proteins with detectable signal from N terminal tagging or C terminal tagging and from at least one terminus (brightest signal), used to assign a protein localisation. **C**. Consensus from N and C terminal tagging, broken down by organelle. For each organelle, N indicates the localisation determined by N terminal tagging of a protein which localised to that organelle by C terminal tagging, and vice versa for C. **D**. Examples of known proteins with known targeting sequences with a mismatch in localisation or lack of signal when tagged at one terminus: BiP, signal peptide disrupted by N terminal tagging. FBPase, glycosomal targeting sequence disrupted by C terminal tagging. NCA2, mitochondrial presequence disrupted by N terminal tagging. Icons in the top right indicate whether an ortholog is present in humans or yeast (*S. cerevisiae* or *S. pombe*). **E**. Euler diagrams of agreement of mass spectrometry-based high confidence proteomes ^24,25,118–123^ with our protein localisations data, with localisations further broken down by organelle substructures for the flagellum and nucleus. **F**. Examples of known paralogous pairs of proteins with one highly expressed in the procyclic form and one in the bloodstream form ^31,124–127^, shown in pairs with the procyclic-specific protein on the left. Both images in each pair are shown with equal contrast.

**Figure S2.**
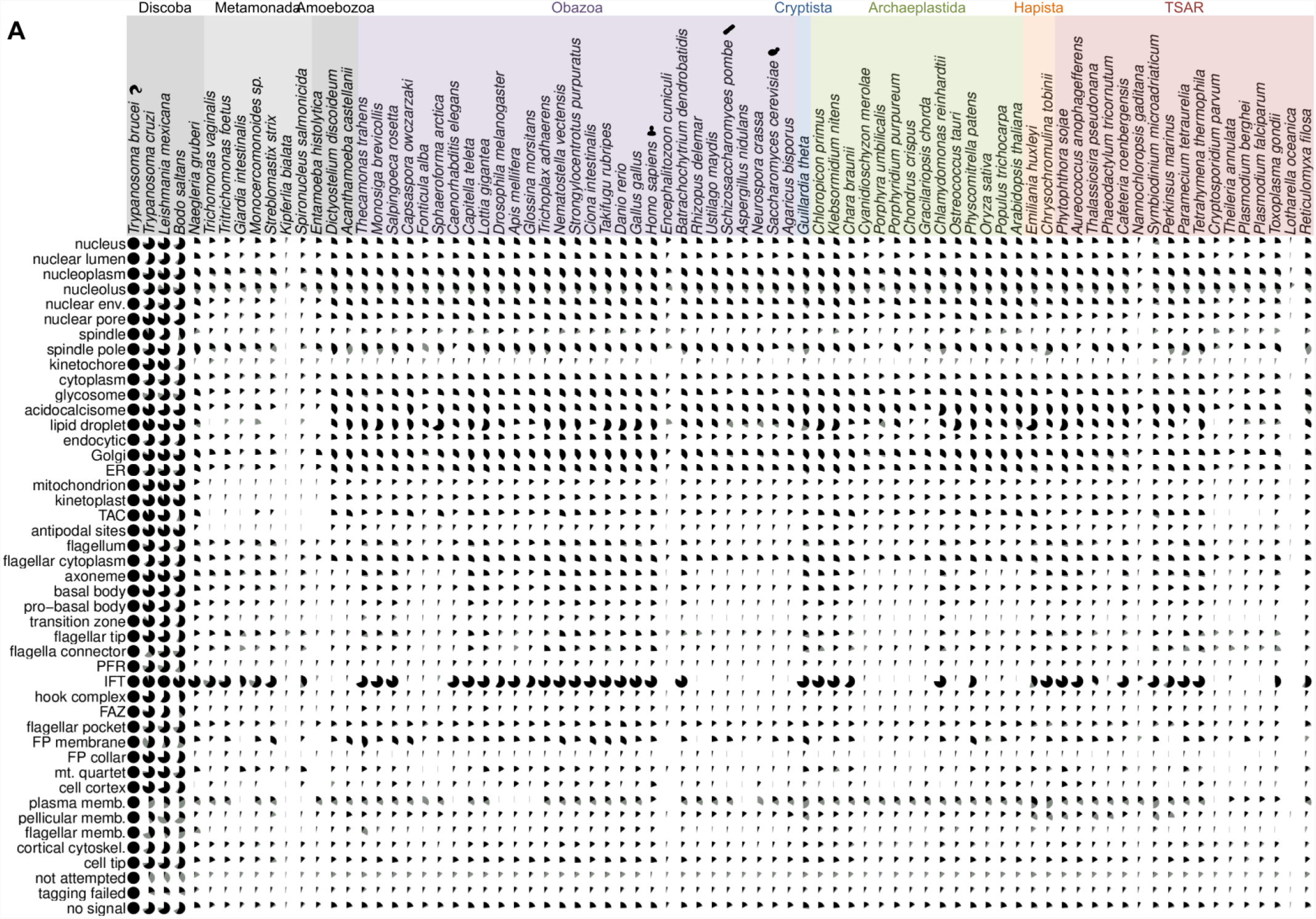
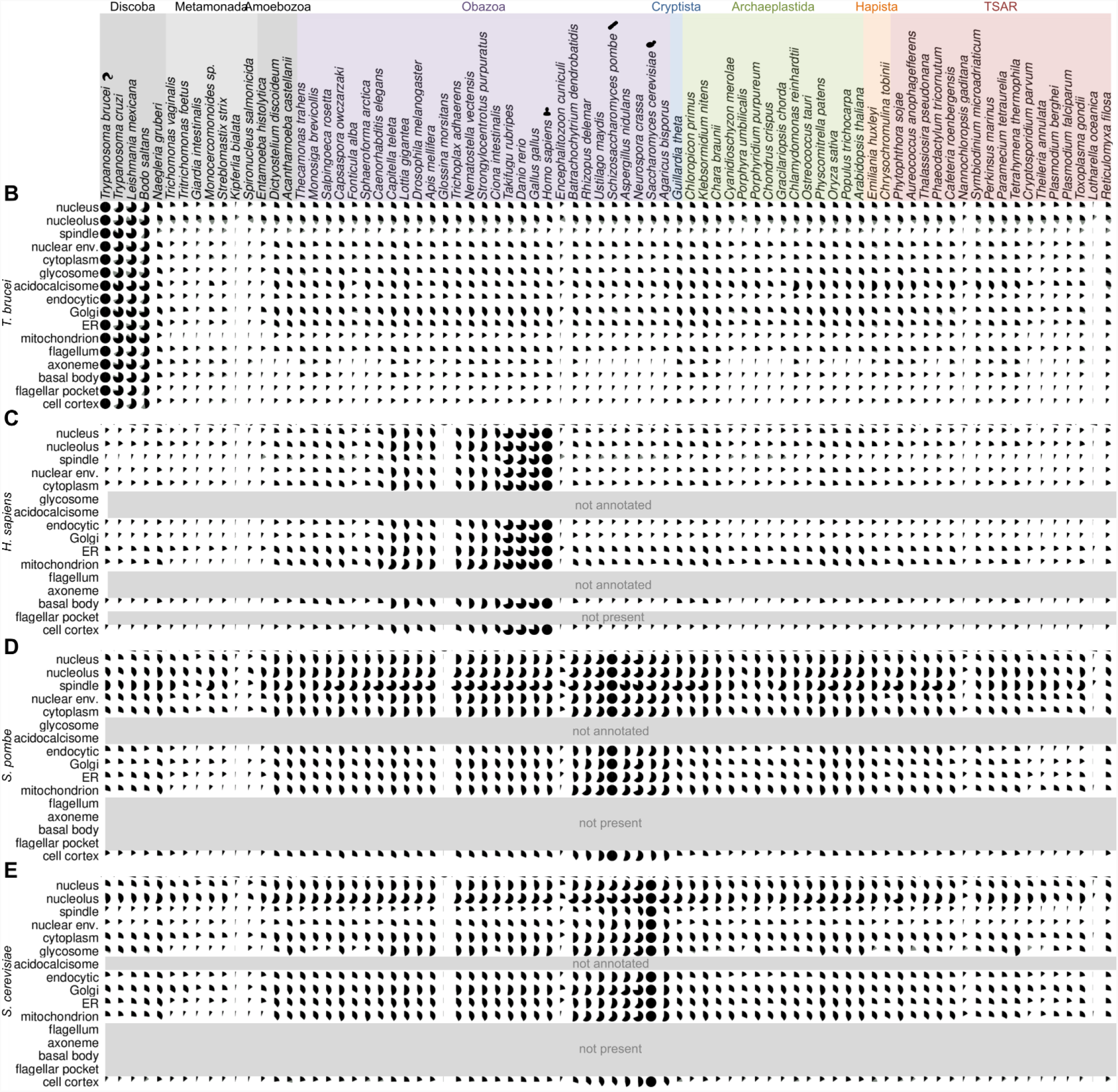
Extended mapping of the universally conserved features of eukaryotic organelles. Extended version of Figure 2. **A**. Presence of orthologs of *T. brucei* proteins, grouped by organelle, across eukaryotic life. Pies represent the proportion of proteins with a reciprocal best BLAST (RBB, black) or not an RBB but at least one orthogroup member (grey) in each species. **B**. As for A, but using a simplified set of localisation annotations. **C-D**. As for B, but plotting orthologs of human proteins (C), *S. pombe* (D) and *S. cerevisiae* (E) proteins, grouped by localisation in their respective genome wide protein localisation projects using a set of terms comparable to B.

**Figure S3.**
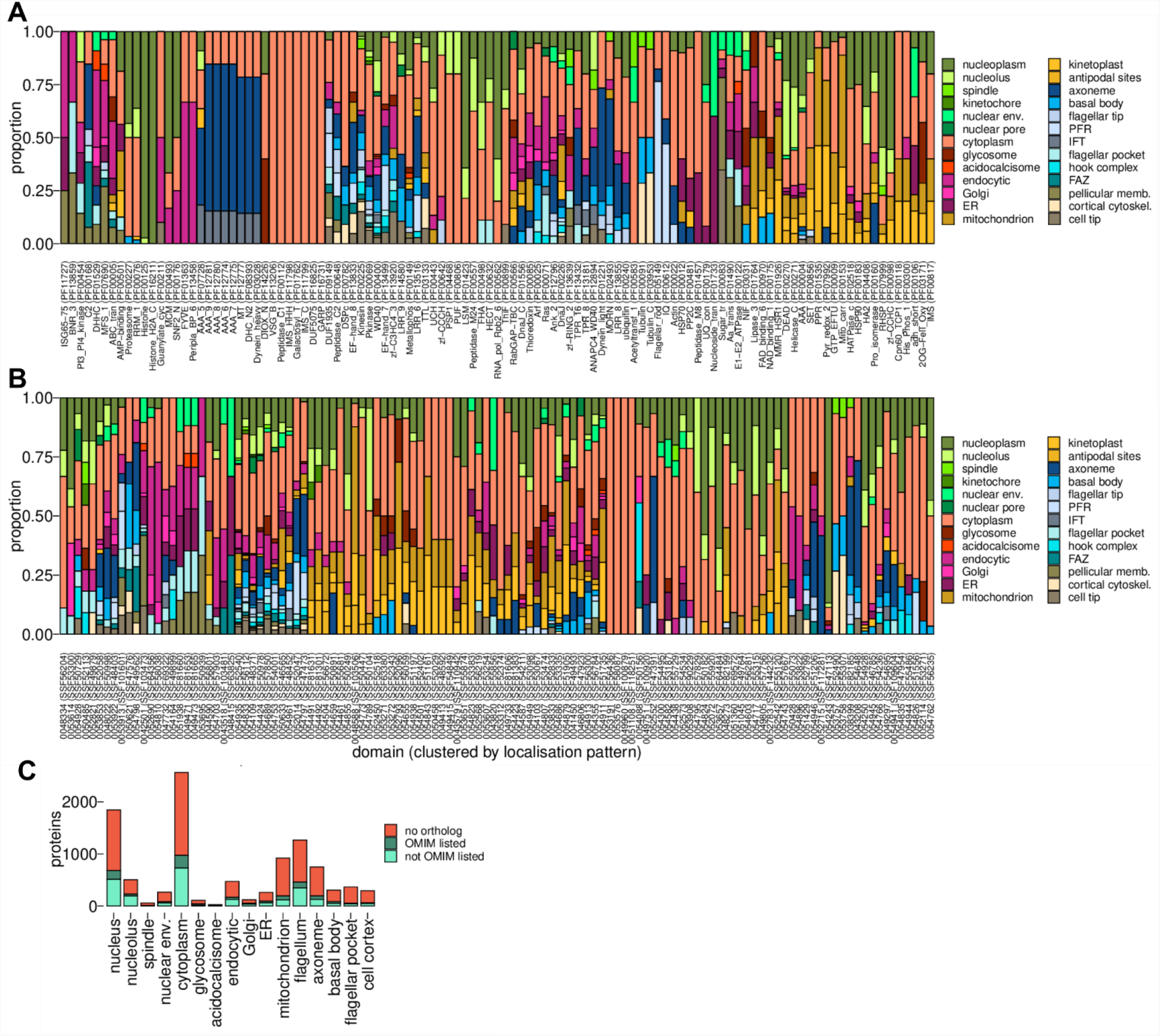
Protein domains and conserved proteins implicated in human genetic disease. **A**. Localisation annotations for all proteins with annotated PFAM domains, grouped by PFAM domain for each domain found in at least 10 proteins. **B**. As for A, except for Superfamily protein domains. **C**. *T. brucei* proteins with a human ortholog, summarised by where the *T. brucei* protein localises and whether the human ortholog is listed as involved in human genetic disease in OMIM.

**Figure S4.**
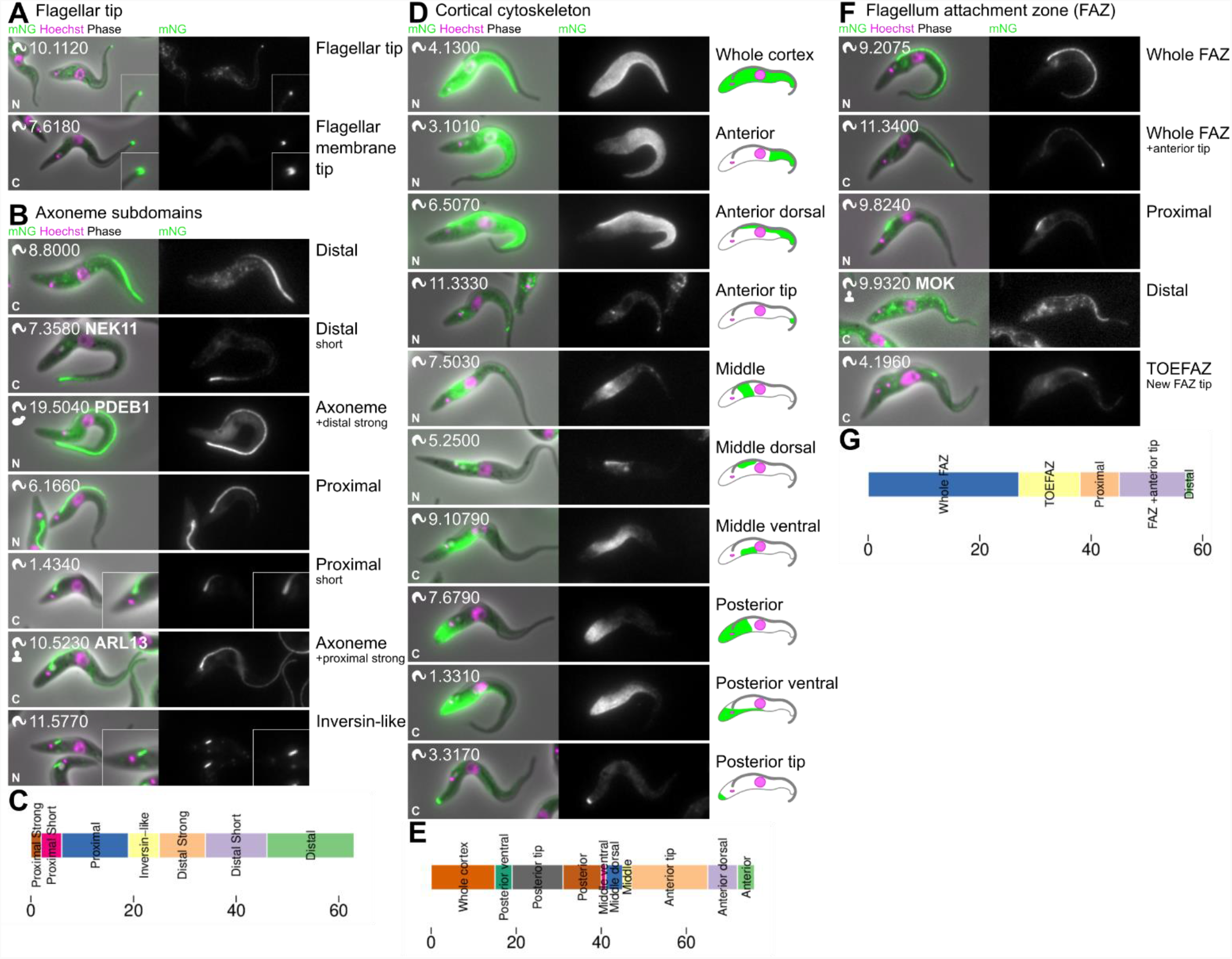
Sub-domains of the flagellar and cellular cytoskeleton implicated in morphogenesis. **A**. Examples of the two major flagellar tip localisations – a small point (axoneme associated) or a horseshoe shape (membrane associated). **B**. Examples of proteins localising to the 7 readily identifiable proximal and distal flagellum, likely axoneme, subdomains. **C**. Number of proteins localising to different proximal and distal domains. **D**. Examples of proteins localising to a range of sub-domains of the microtubule-based cortical cytoskeleton. At least 10 sub-domains, 7 novel, exist. **E**. Number of proteins localising to each cortical cytoskeleton sub-domain. **F**. Examples of proteins localising to a range of sub-domains of the flagellum attachment zone (FAZ) which is a specialised seam in the cortical cytoskeleton required for lateral attachment of the flagellum. At least 5 sub-domains exist. The anterior cell tip and distal FAZ tip appear synonymous. Some FAZ proteins localise only to the tip of the extending FAZ (TOEFAZ). **G**. Number of proteins localising to each FAZ sub-domain

**Figure S5.**
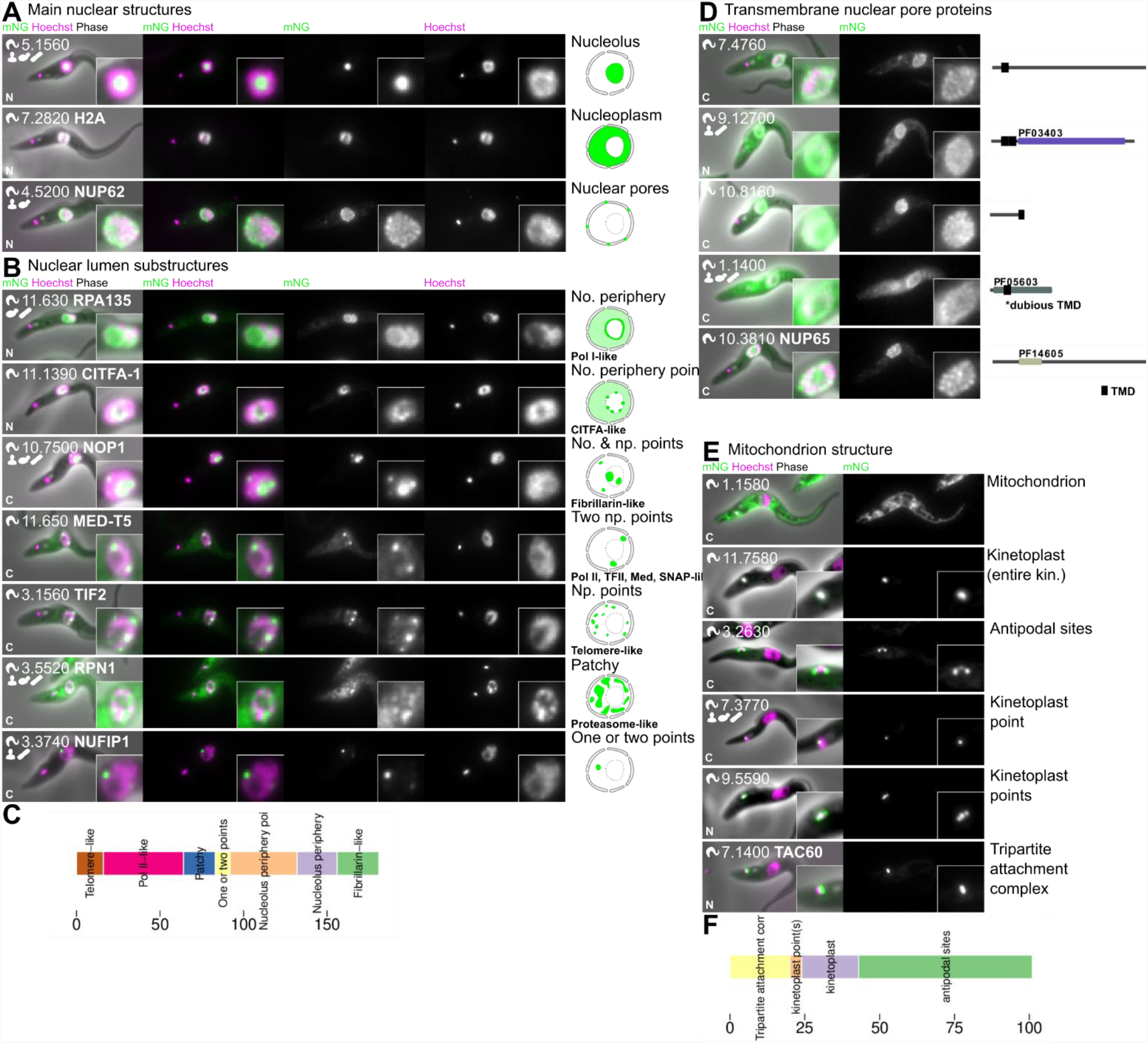
Non-membrane bound complexity of the nuclear and mitochondrial DNA compartments. **A**. Examples of proteins localising to major nuclear subcompartments: the nucleolus, nucleoplasm and the nuclear pores. **B**. Examples of characterised proteins localising to a range of sub-domains within the nuclear lumen. 7 characteristic localisation patterns are visible. Likely functions in Pol I, Pol II, nucleolar (fibrillarin), telomere and nuclear proteasome function are ascribable to other proteins with a similar localisation. **C**. Number of proteins localising to each nuclear lumen sub-compartment (excluding simple nucleolar and nucleoplasmic localisations). **D**. All five proteins localising to the nuclear pores with predicted transmembrane domains, shown next to a cartoon representation of the protein domain structure. All except NUP65 (Tb927.10.3810) are novel. **E**. Examples of proteins localising to different mitochondrion and kinetoplast structures **F**. Number of proteins localising to each kinetoplast associated structure.

**Figure S6.**
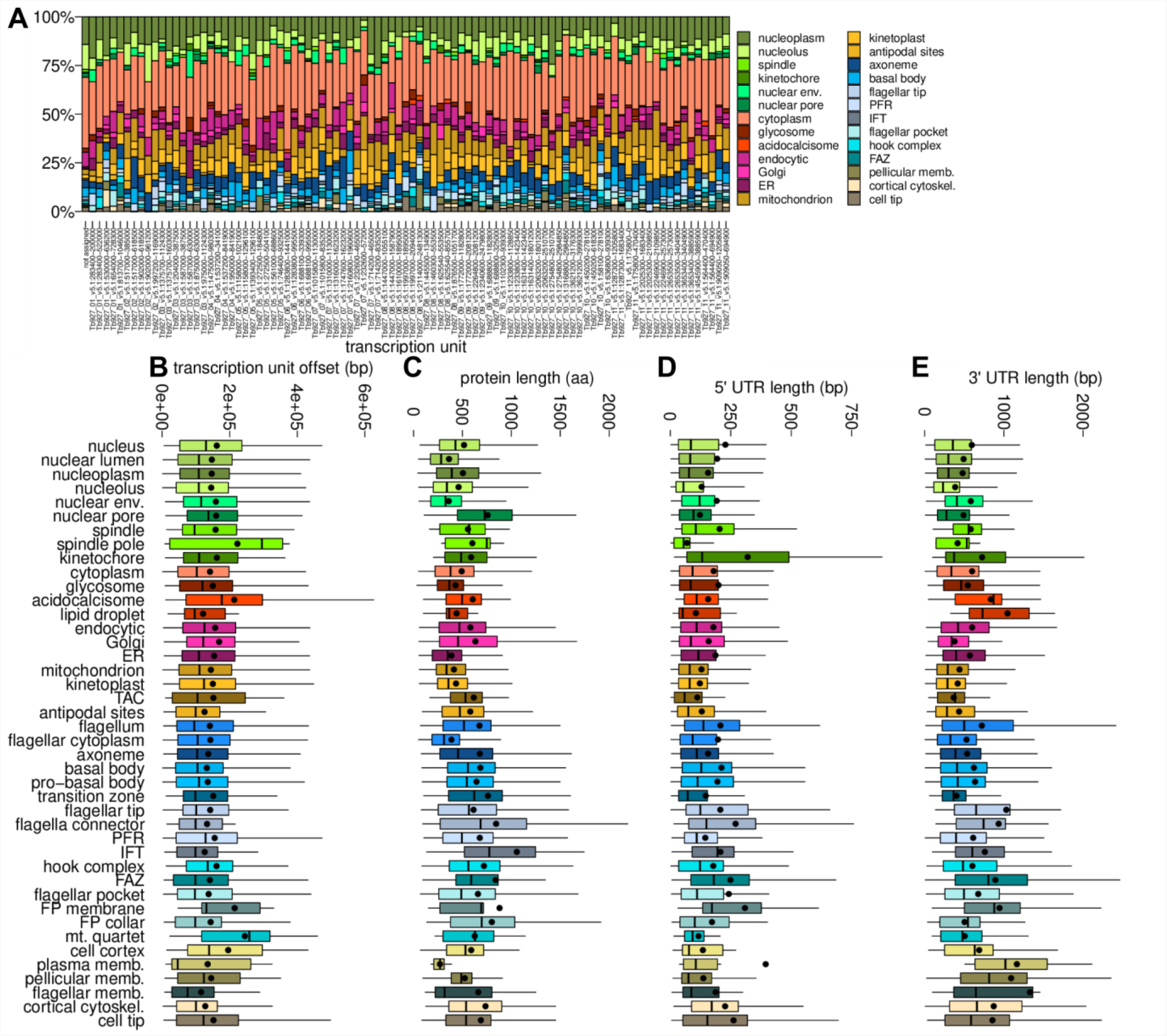
Gene transcription units do not reflect grouping in protein localisation. **A)** Localisation terms for proteins encoded by each transcription unit. **B-E)** Gene properties, broken down by protein localisation. Box represents the median and interquartile range, whiskers represent the 5^th^ and 95^th^ percentile and the dot represents the mean. B) Offset in base pairs of the start of the gene from the start of the transcription unit. C) Predicted protein size in amino acids. D) 5′ UTR length. E) 3′ UTR length.

Table S1. Localisation annotation ontology, including GO terms

Table S2. Localisation modifier ontology

Table S3. The localisation dataset

Table S4. Genomes and accessions

Table S5. Human and yeast localisation annotation term mappings

